# Predicting Species-Area Relationships in Fragmented Forests

**DOI:** 10.64898/2026.09.30.755608

**Authors:** Monique de Jager, Finn J. van Wordragen, Fleur Vrouwenvelder-Slegers, Edwin T. Pos

**Author notes:** **Corresponding author:** Monique de Jager.

## Abstract

Habitat destruction leads to a severe decline in biodiversity. Ascertaining effects of habitat loss afterwards is regularly done, whereas predicting the effects of different habitat loss scenarios on biodiversity beforehand is much preferred but as of yet rarely possible. Such predictions would help inform management decisions to prevent biodiversity loss in the first place, or at least mitigate effects if necessary, by providing handles to set priorities in making trade-offs.

To this end, we developed and applied an adapted version of the information-theoretical framework of Maximum Entropy Theory in Ecology (METE) to predict species-area relationships in tropical forest landscapes undergoing habitat loss and fragmentation. Species-area relationships provide information about biodiversity and can, in undisturbed systems, be predicted using METE, based on state variables such as the total number of individuals, the total number of species, and the total area size. In disturbed systems, however, it is established that METE’s predictions are off. Using an individual-based model, we explore how METE’s state variables should be adjusted, based on the landscape characteristics habitat cover and degree of habitat fragmentation, to restore its accuracy of species-area relationship predictions in disturbed environments.

We validate our adjusted METE predictions with empirical data. Our adjusted METE predictions demonstrate the severity of fragmentation’s impact on biodiversity, relative to what could be expected in a pristine, continuous environment. With this new method, we can estimate the immediate effects of habitat loss on biodiversity, as well as its long-term effects.

## 1. INTRODUCTION

Maintaining high biodiversity levels is of key importance in keeping ecosystems healthy and resilient, especially in landscapes undergoing intensive anthropogenic disturbances (Loreau et al. 2001). Although biodiversity loss accelerates the decline in ecosystem productivity, human activities like resource extraction, land use changes, and habitat destruction continue to significantly reduce biodiversity.

It is well known that habitat destruction leads to severe biodiversity decline. In addition, the specific spatial configuration of habitat loss may affect its impact (Quinn and Harrison 1988; Tscharntke et al. 2002; Campos et al. 2012; Hanski et al. 2013; Haddad et al. 2015; Keil et al. 2015; Fahrig 2017; Seibold et al. 2017; Chisholm et al. 2018; Fahrig et al. 2019). The spatial configuration of habitat loss is often referred to as habitat fragmentation *per se* (Fahrig 2003; May et al. 2019), which we hereafter will call habitat fragmentation. The effects of habitat fragmentation depend on the spatial and temporal scales at which biodiversity loss is measured (de Jager and Pos 2026). For example, at the landscape scale, habitat loss initially causes the greatest biodiversity decline in habitats with low fragmentation. Over time, however, biodiversity loss intensifies in highly fragmented environments as community dynamics stabilize, eventually surpassing that observed in less fragmented landscapes (de Jager and Pos 2026).

Quantifying the effects of habitat loss and fragmentation on species richness is essential for assessing biodiversity impacts and informing conservation decisions. Predicting the effects of alternative habitat-loss scenarios before they occur would therefore be particularly valuable for management. Such predictions have been attempted using spatial simulations (e.g. Hubbell et al. 2008; Campos et al. 2013) and modified species–area relationship (SAR) models (e.g. Pereira and Daily 2006; Hanski et al. 2013; Kuipers et al. 2021). Although these approaches can provide valuable predictions of biodiversity responses to habitat loss, their application often depends on detailed ecological data, model-specific parameterization, and specialized analytical expertise. This can limit their transferability and practical use across landscapes and management (Hanski et al. 2013; Larrey-Lassalle, Loiseau, et al. 2018; Larrey-Lassalle, Esnouf, et al. 2018; Chisholm et al. 2018; Kuipers et al. 2021). Moreover, a general and readily applicable approach for predicting the longer-term biodiversity consequences of alternative habitat-loss and fragmentation scenarios remains lacking.

Here, we develop such an approach by adapting the Maximum Entropy Theory in Ecology (METE) framework (Harte 2011; Harte and Newman 2014). METE is particularly useful in this context because it provides an analytically tractable and parsimonious link between a limited set of system-level state variables and biodiversity patterns. Previous applications have shown that METE can generate accurate SAR predictions without requiring the full species-specific parameterization and computational complexity of an individual-based model for every new scenario (Harte 2011; Harte and Newman 2014). It is well-known, however, that METE is limited in prediction accuracy for systems that are in flux, i.e. for which the aggregated measures of state variables like the total number of species or individuals are changing (Newman et al. 2020; A. J. Rominger et al. 2016; Carey et al. 2006; Kempton and Taylor 1974; Supp et al. 2012). To restore accuracy, we use individual-based model simulations to determine how METE’s state variables should be adjusted under different landscape configurations, and validate the resulting predictions against empirical data. We show that habitat cover and the degree of habitat fragmentation can be used to predict SARs in tropical forest landscapes undergoing habitat loss and fragmentation, thereby providing a practical framework for estimating biodiversity loss under alternative habitat-loss scenarios. In addition, our adjusted framework also provides an interpretable baseline for quantifying biodiversity loss relative to that expected in an undisturbed, continuous landscape.

## 2. METHODS

Species-Area Relationships (SARs) describe the distribution of species richness over subcommunities within a landscape, specifically the relationship between area size (*A*) and the number of species present in an area of that size (*S*) (Connor and McCoy 1979). Macroecological probability distributions, such as the SAR, can be derived using METE from only some observed (macroscopic or aggregated) state variables of the total community (Harte 2011; Harte and Newman 2014). In the original framework, these state variables are (i) the total ecosystem area (*A*_*0*_), (ii) the total number of individuals (*N*_*0*_), (iii) the total number of species (*S*_*0*_), and, optionally, (iv) the total metabolic energy rate (*E*_*0*_; all within *A*_*0*_). These state variables impose constraints on the probability distribution of interest (e.g., the SAR), ensuring that the derived distribution matches their observed values. Otherwise, the derived probability distribution is maximally uninformed, distributing probability as evenly as possible given the imposed constraints.

The only explicit scaling relationship assumed by METE is how the abundance of a species scales with area size *A*, where *A* is a subarea of the total area *A*_0_. This scaling relationship is evident in the constraint that METE imposes on the Spatial Structure Function Π(*n*|*A, n*_0_, *A*_0_):

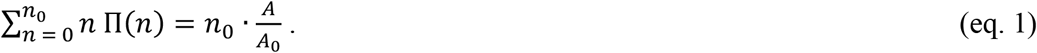

Here, Π(*n*) is the probability that *n* individuals of a species are found in an area of size *A*, given that the species has *n*_0_ individuals in the total area *A*_0_. This constraint enforces that the expected abundance in area *A* equals 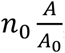, i.e., abundances scale linearly with area (Brummer and Newman 2019), thereby implying that individuals of each species are distributed continuously and uniformly across the landscape. Yet, given the often-observed clustered distributions of species, this assumption can be violated. Moreover, fragmentation can reduce species richness beyond the assumed scaling relationship. The exact amount of species loss depends, in part, on the geometric form of habitat destruction (de Jager and Pos 2026), arguing for a new perspective when applying the METE framework to fragmented landscapes.

In the case of the violated assumption of the scaling relationship, METE is likely to fail for systems under disturbance, as has been shown earlier (Brush et al. 2022; Franzman et al. 2021; Newman et al. 2020). Here, however, we revise the model by altering its state variables in a backwards approach, by exploring how much the state variables *N*_*0*_ and *S*_*0*_ should be modified to provide accurate estimations of SAR using METE in a wide range of differently fragmented landscapes. By adjusting constraints that deviate from the expected scaling relationship, we show improved predictions under potential future habitat-loss scenarios. In addition, using this approach, we can estimate the immediate effects of habitat loss on biodiversity loss, as well as the long-term effects (i.e. after community dynamics have stabilized). The adjusted METE predictions inform us of the severity of fragmentation’s impact on biodiversity, relative to what could be expected in a pristine, continuous environment. To obtain adjusted state variables that allow METE’s SAR predictions to be more accurate in fragmented environments, we simulated community dynamics in Amazonian rainforests using an individual-based model, and matched the SARs retrieved from these simulations with the best fitting METE predictions.

### 2.1 An individual-based model of Amazonian tree communities

We simulated community dynamics of an Amazonian tree (meta)community using a semi spatially-explicit, near-neutral, individual-based model (de Jager and Pos 2026). The modelled environment consists of a 2D-lattice containing 2025 patches (a 45 by 45 grid) categorically subdivided into habitable or uninhabitable patches. A habitable patch holds a subcommunity of 1,000 individuals; no individuals are present at uninhabitable patches. Assuming a tree density of 400 trees ha^-1^ (ter Steege et al. 2003), patch size roughly equals 2.5 ha and the total simulated area is approximately 50 km^2^.

We first created a stable community (reflecting a pristine, continuous environment) to initialize our simulations: we filled the subcommunities with a total of 2,025,000 individuals that were randomly drawn from a species-pool consisting of *S*_*max*_ species. Second, we iterated community dynamics (following steps outlined below) 10,000 times to achieve a stable community.

Each iteration of community dynamics involves the following five steps: (1) Patches are selected in random order. (2) Within the selected patch, individuals present at the beginning of the iteration are selected in random order. (3) The selected individual produces one offspring and disperses it. Seed dispersal occurs in a random direction, across a distance (*d*, in patches) that is randomly drawn from a bounded pareto distribution:

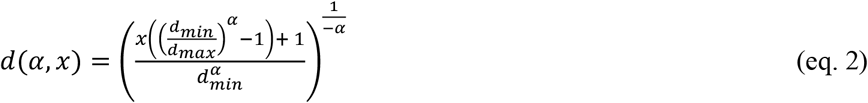

Here, *d*_*min*_ and *d*_*max*_ are the minimum and maximum dispersal distances (*d*_*min*_ = 0.01 and *d*_*max*_ = 6.5 patches), *x* is a random number drawn from a uniform distribution (0 ≤ *x* < 1), and *α* is a scaling exponent. This scaling exponent depends on the species identification *s* (*α = s/S*_*max*_, 0 < α ≤ 1): species with a large identification number disperse more locally and species with a small identification number disperse over a longer range. Within the dispersed-to patch, a random individual is chosen to be replaced by the dispersed seed in the next step. (4) After all individuals have reproduced and dispersed, the dispersed seeds replace the individuals that previously occupied those sites. If an individual was selected multiple times for replacement, the seed that dispersed last replaces it. If, after all individuals have reproduced, some individuals remain unreplaced, they stay at their sites for the next iteration. (5) Individuals are replaced by offspring from a randomly selected species through weighted random selection based on the species’ dispersal capacity. Species with larger dispersal capacities have higher probabilities of being introduced into subcommunities from the larger metacommunity with probability *P*_*meta*_, representing vagrant dispersal from a larger species pool containing all *S*_*max*_ species (in equal abundances).

### 2.2 Model calibration

Before introducing habitat loss and fragmentation to the model simulations, we calibrated the model’s parameters: (1) the maximum dispersal distance *d*_*max*_, (2) the probability that an individual is replaced by a seed from the metacommunity *P*_*meta*_, and (3) the number of species available in the metacommunity *S*_*max*_. First, we ran simulations with 300 different combinations of *d*_*max*_ (4.5 – 6.5, with 0.5 increments), *P*_*meta*_ (0.001 – 0.002, with 0.0002 increments), and *S*_*max*_ (5,500 ≤ *S*_*max*_ ≤ 10,000, with increments of 500 species), using a full orthogonal sampling of the 3-dimensional parameter combination matrix. We calculated a goodness-of-fit (GOF) by fitting our simulation results to two empirical datasets (4531 species in 10.72 km^2^ in 2013, and 5027 tree species in 20.24 km^2^ in 2019; ter Steege et al. 2020):

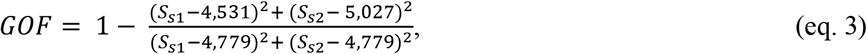

where *S*_*s1*_ and *S*_*s2*_ are the number of species observed in our simulations within an area of 10.72 and 20.24 km^2^, respectively. Values of *GOF* close to 1 indicate that the simulation results are close to empirical observations. We selected the parameter value combination that resulted in the highest GOF to be used in the main simulations. Because this parameter value combination was on the edge of the parameter combination matrix, we performed 68 additional simulations, thereby expanding the parameter value ranges to 4.5 ≤ *d*_*max*_ ≤ 7.5, 0.001 ≤ P_*meta*_ ≤ 0.0024, and 4,500 ≤ *S*_*max*_ ≤ 10,000. The best fitting parameter value combination was *d*_*max*_ = 6.5, *P*_*meta*_ = 0.002, and *S*_*max*_ = 5500. These values were subsequently used in all our other simulations.

### 2.3 Introducing habitat loss and fragmentation

We generated environments with different levels of habitat loss and fragmentation using weighted random patch selection, starting from a random initial habitat patch. We successively added new habitat patches, until the correct habitat cover for that specific environment was reached. Patches were selected using a weighted random selection. Weights (*W*_*i*_) were calculated from distances to already selected habitat patches using a 2-D pareto distribution:

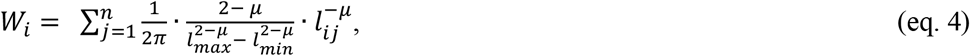

where *l*_*ij*_ is the Euclidean distance between non-habitat patch *i* and habitat patch *j, n* is the number of habitat patches, *l*_*min*_ and *l*_*max*_ are the minimum and maximum distances to consider (*l*_*min*_ = 1, *l*_*max*_ = 50 patches), and *µ* is the scaling exponent (*µ* ≥ 1). By altering the scaling exponent *µ* of this distribution, landscapes with different degrees of fragmentation can be generated. *µ* = 1 results in the most fragmented environments; habitat patches become more clustered with increasing *µ*.

With the calibrated model, we ran simulations with 90 different combinations of habitat loss (0 – 90% habitat loss, with 10% increments) and habitat fragmentation levels (1 ≤ *µ* ≤ 5, with 0.5 increments), again using a full orthogonal sampling of the 2-dimensional parameter combination matrix. Each simulation was run 5 times to account for the stochastic nature of the model.

Directly after habitat loss as well as after 10,000 iterations of community dynamics, we recorded the number of individuals per species per habitat patch, the percentage habitat cover, and Moran’s I. Moran’s I is an indicator of spatial heterogeneity; low Moran’s I values correspond to high fragmentation, while high Moran’s I values correspond to high spatial clustering of habitat patches. Habitat cover and Moran’s I can be easily estimated from satellite images and are therefore useful parameters for predicting the effects of habitat loss and fragmentation on species richness. Using the recorded subcommunity compositions, we calculated the corresponding SARs per simulation, which we refer to as the observed SARs.

### 2.4 Calculating SAR

For a range of area sizes (*A*_*i*_ = 0.005, 0.01, 0.02, 0.04, 0.08, 0.16, 0.32, 0.64, 1.28, 2.56, 5.12, 10.24, and 20.48 ha), we estimated the number of unique species as the average of 100 draws from all 2,025,000 potential individuals, including empty spaces, randomly selecting an equivalent number of individuals that would fit the area, assuming each occupies .0025 ha (*n*_i_ = 2, 4, 8, 16, 32, 64, 128, 256, 512, 1024, 2048, 4096, and 8192, respectively; ter Steege et al. 2017).

### 2.5 Predicting SAR with METE

Meanwhile, we generated multiple SARs with METE by systematically varying *N*_0_ and *S*_0_. We generated METE predictions using the *meteSAR* function from the R-package *meteR* (using R version 4.5.0, Rominger and Merow 2017). This function requires a minimum area size (*A*_*min*_ = 0.005 ha), a maximum area size (*A*_*0*_ = 41 ha), the total number of species present within the total area (*S*_*0*_), and the total number of individuals (*N*_*0*_). We calculated METE predictions of the SAR for 5940 combinations of *S*_*0*_ and *N*_*0*_ (200 values of *S*_*0*_, 55 ≤ *S*_*o*_ ≤ 11,000 and 40 values of *N*_*0*_, 512 ≤ *N*_*0*_ ≤ 20,480, with increments of 55 and 512, respectively).

In METE, the SAR is derived from its predictions of the Spatial Structure Function (SSF) and the species abundance distribution (SAD). The SSF is a species-level prediction of how individuals cluster in space. It is the probability that *n* individuals of a species are in a cell of area *A*, given that the species has *N*_0_ individuals in *A*_0_:

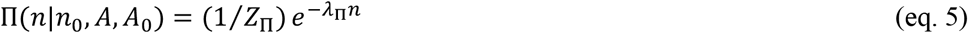

with partition function 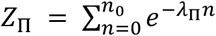 normalizing the probability distribution. The SAD gives the probability that a species has abundance *n* within a community of *N*_0_ individuals and *S*_0_ species:

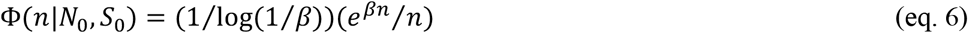

METE calculates the SSF and SAD by maximizing Shannon Entropy under constraints, with normalization always included to ensure probabilities sum to one. For the SSF, an additional constraint is imposed that the abundance of a species increases proportionally with the area size:

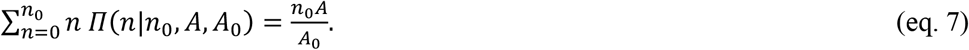

For the SAD, two extra constraints are imposed: the average abundance per species, ⟨*n*⟩ = *N*_0_⁄*S*_0_, and the average metabolic energy per species, ⟨*n*ε⟩ = *N*_0_⁄*S*_0_. Here, ε and *E*_0_ represent the individual and total metabolic rates in the community, respectively. Since we do not provide metabolic data, *E*_0_ defaults to a standard value. METE determines the parameters (Lagrange multipliers) that maximize entropy while meeting the constraints. The variables *λ*_Π_ in eq. 5 and *β* in eq. 6 depend on these multipliers.

METE’s SAR (eq. 8) combines the SSF and the SAD. The SAD supplies the probability that a species has abundance *n* in the full area *A*_0_; the SSF then gives the probability that a species present with *n* individuals in *A*_0_ has no individuals in a cell of area *A* (*Π*(0|*n, A, A*_0_)). Its complement is the probability that the species is detected somewhere in the smaller area. Averaging this detection probability over the SAD and scaling by *S*_0_ yields the expected number of species in area *A* (i.e., the SAR):

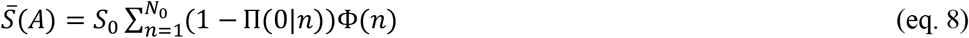

For a full overview of functions in the METE framework, we refer to the original publications (Rominger and Merow 2017; Brummer and Newman 2019; Harte et al. 2008).

### 2.6 Comparing SARs from simulations and METE predictions

For each combination of SAR observed in a simulation and predicted by METE, we calculated a Goodness-Of-Fit (*GOF*_*METE*_):

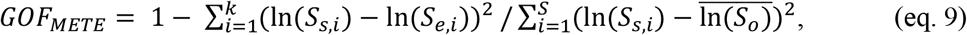

where *k* is the number of datapoints to compare (*k* = 13, at *A*_*i*_ = 0.005, 0.01, 0.02, 0.04, 0.08, 0.16, 0.32, 0.64, 1.28, 2.56, 5.12, 10.24, and 20.48 ha), *S*_*s,i*_ and *S*_*e,i*_ are the number of species observed in simulations and expected by METE estimation for area *A*_*i*_, respectively, and 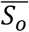 is the average number of species.

GOF-values close to 1 indicate that the prediction by METE of the SAR is highly comparable and, hence, the metacommunity appears to be in a state that would adhere to what is expected based on the macroscopic constraints. For each combination of habitat cover, degree of clustering (in terms of Moran’s I), and time since the disturbance (immediately after habitat loss or after community dynamics have stabilized), we selected and recorded the (*N*_*0*_, *S*_*0*_)-combination that resulted in the best GOF, giving us the adjusted state variables 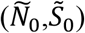. Because the relations between the adjusted state variables and the environmental parameters habitat cover and habitat clustering were nonlinear, we did not fit this relation into a simple equation, but rather directly selected the corresponding 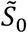 from the data frame to estimate SARs with METE in areas undergoing habitat destruction.

### 2.7 Validating our Predictions of Species-Area Relationships

To predict the SAR of a real-world area undergoing habitat destruction, we can adjust its observed state variables to the optimal values from simulations that best match the area’s degree of habitat cover and spatial clustering of habitat patches. As an example, we show how observed SARs in fragmented environments are fitted with our method, using the data from the LandFrag dataset (Gonçalves-Souza et al. 2025). We selected six datasets from studies performed after 2014 that held >500 observations each (Magnago et al. 2015; Pinho et al. 2021; Safar et al. 2022; Santos et al. 2007). Each dataset consists of 0.1 ha plots (except Magnago et al. (2015), where plot size was 0.01 ha) in both continuous forest areas and in fragmented environments. Per dataset, we divided the different plots in groups, selecting on proximity and forest type, resulting in 9 study areas (Fig. 1). All study areas were located in the Atlantic Forest area (Fig. 1). Over the last decades, the Atlantic Forest has lost 88% of its native forest cover due to exploitation, agriculture, and urbanization (Fundacao 2022). Between 2020 and 2021, deforestation rates in this area increased with 66% (Fundacao 2022). The vegetation at the sites is classified as Lowland Rainforest or Coastal/Tertiary Tableland forest (Magnago et al. 2015; Pinho et al. 2021; Safar et al. 2022; Santos et al. 2007). The study areas differ in native forest cover, matrix quality, overall forest cover, and degree of forest fragmentation (Fig. 2).

**Figure 1.**
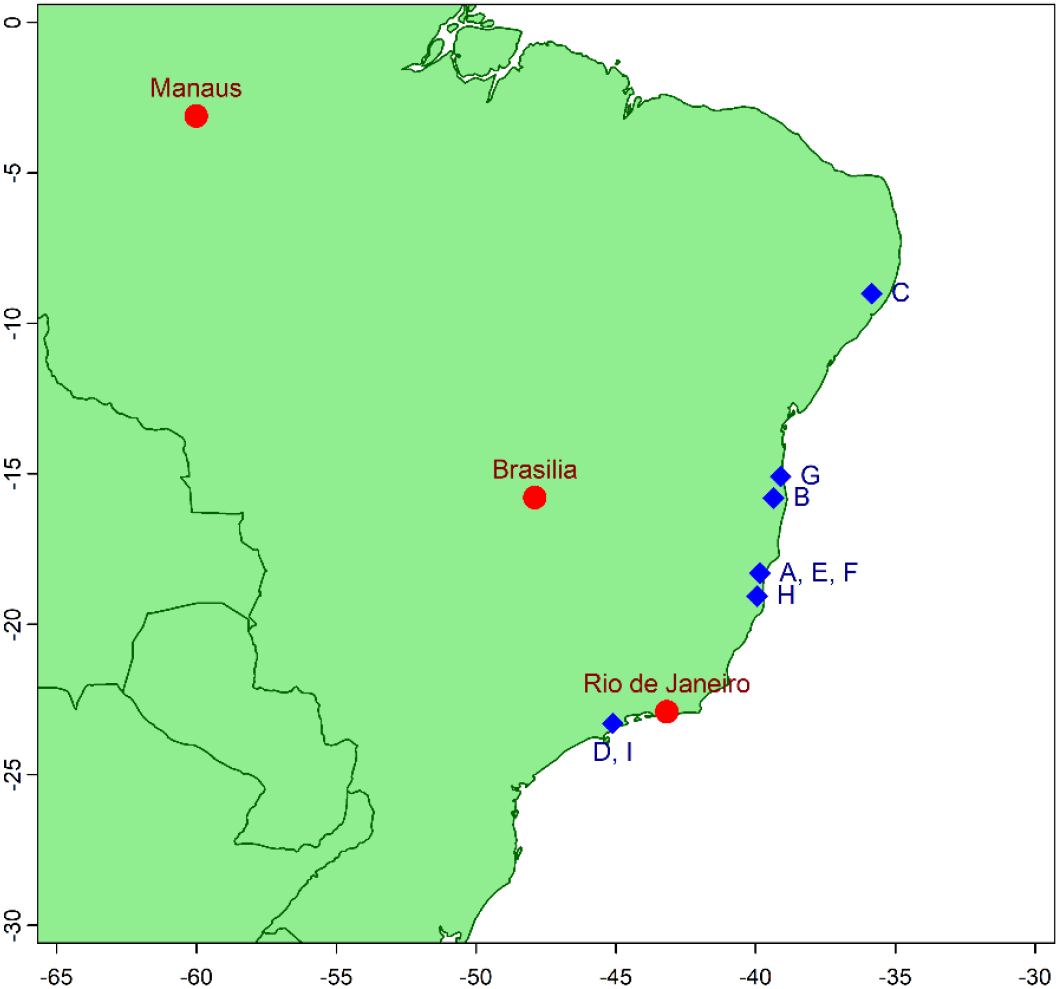
Locations of the nine selected study sites. (**A, E, F**) Safar et al. (2022), (**B, D, G, I**) Pinho et al. (2021), (**C**) Santos et al. (2007), and (**H**) Magnago et al. (2015).

**Figure 2.**
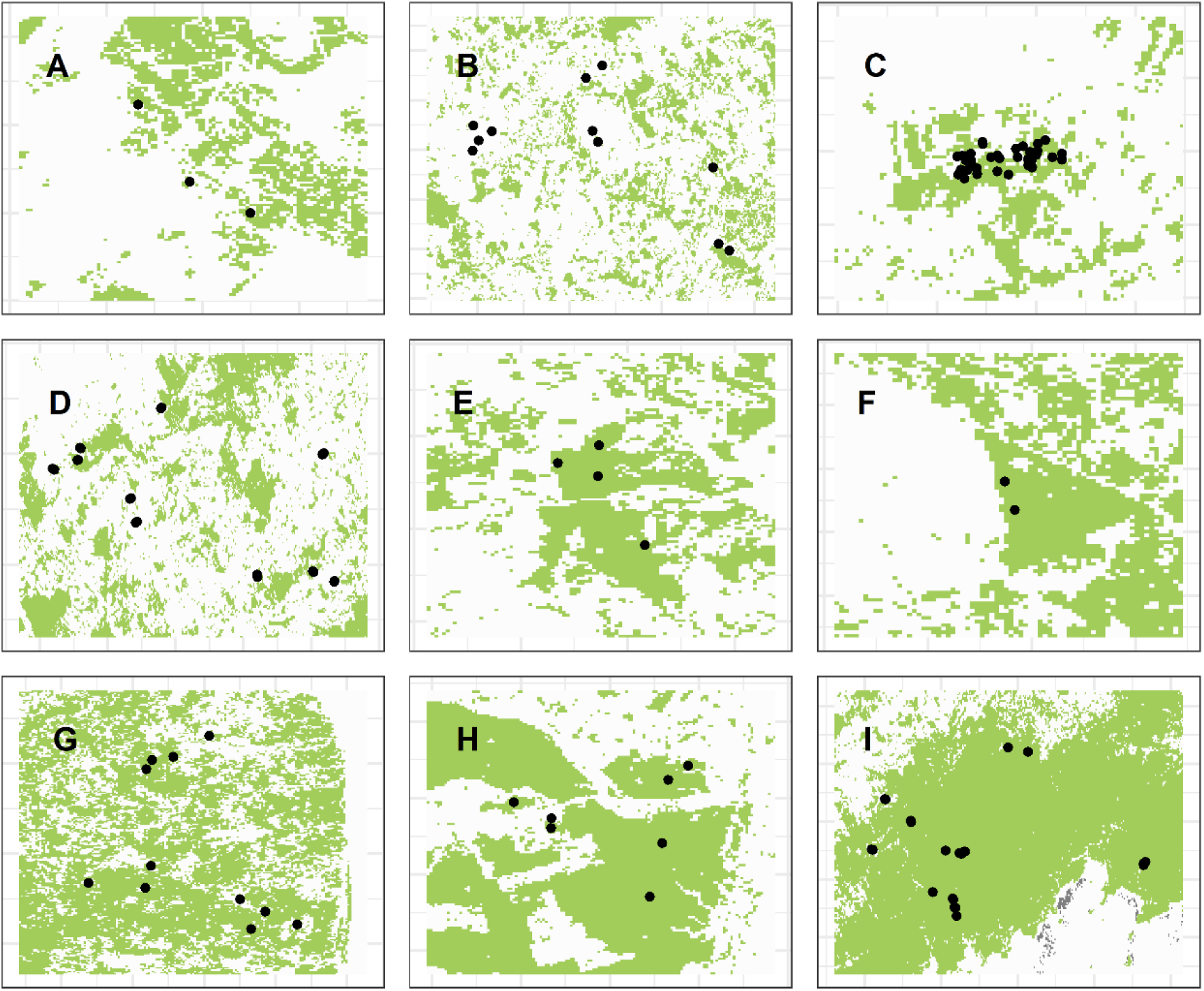
Locations of the different plots selected from the LandFrag dataset, to use as an example of our SAR estimation using the adjusted METE calculations. (**A, E, F**) Safar et al. (2022), (**B, D, G, I**) Pinho et al. (2021), (**C**) Santos et al. (2007), and (**H**) Magnago et al. (2015). Area sizes differ between the study areas, see table 1 for more information.

Per study area, we retrieved Landsat 8 band 4 and 5 data from Google Earth Engine (Gorelick et al. 2017) to calculate the Normalized Difference Vegetation Index (NDVI). Resolution was set at 158 × 158 m, corresponding to the resolution of our model simulations. Otherwise, habitat fragmentation would be measured at a different spatial scale, which would give different results. Per study area, we selected the area that contained all plots, with an additional outer edge of 0.05 longitude and 0.05 latitude. The area sizes ranged between 142 and 1643 km^2^, with an average area size of 743 km^2^ (Table 1). We used an NDVI threshold value (NDVI > 0.8) to create a binary map of forest and non-forest cells (Fig. 2). NDVI values ≤ 0.8 were counted as non-forest area, and values > 0.8 as forest areas (Nath et al. 2024). Using this binary map, we calculated the percentage forest cover and the degree of clustering of forest cells (in terms of Moran’s I; table 1).

**Table 1.** Total number of observations (n_sampled_), number of plots, area size (in km^2^), % forest cover (c), and Moran’s I per study area.

|  | Study | # Observations | # Plots | Area size (km<sup>2</sup>) | % Cover | Moran's I |
| --- | --- | --- | --- | --- | --- | --- |
| A | Safar et al. (2022) | 295 | 30 | 294 | 25 | 0.61 |
| B | Pinho et al. (2021) | 459 | 11 | 1,348 | 24 | 0.57 |
| C | Santos et al. (2007) | 2,983 | 37 | 205 | 19 | 0.60 |
| D | Pinho et al. (2021) | 3,399 | 20 | 1,642 | 26 | 0.68 |
| E | Safar et al. (2022) | 556 | 40 | 251 | 36 | 0.61 |
| F | Safar et al. (2022) | 291 | 20 | 142 | 41 | 0.62 |
| G | Pinho et al. (2021) | 1,267 | 11 | 960 | 57 | 0.55 |
| H | Magnago et al. (2015) | 1,199 | 70 | 455 | 53 | 0.81 |
| I | Pinho et al. (2021) | 15,083 | 23 | 1,392 | 71 | 0.82 |

Using the percentage habitat cover ± 5% and the degree of spatial clustering ± 0.05, we selected the eight corresponding adjusted state variables; the four closest values of *Ñ*_0_ and 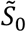 immediately after habitat loss and the four closest values of *Ñ*_0_ and 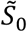 after community dynamics have stabilized. With these adjusted state variables as input for METE, we created two regions in the species-area plot: one for the SAR immediately after habitat loss has occurred, and one for the SAR after community dynamics have stabilized.

We estimate the empirical SAR using the plot data, which contains the species name of each individual tree within the selected plots. We used the same method to obtain the empirical SAR as we used with the simulation data. In line with this method, we added samples from uninhabitable, empty locations. We calculated the area size of these empty locations per study area as *n*_*sampled*_ / *c*-*n*_*sampled*_, where *n*_*sampled*_ is the total number of individuals sampled and *c* is the estimated forest cover (Table 1). We added these empty locations in our random sampling to estimate the SAR. The empirical SARs were then plotted on top of the SAR areas predicted by METE.

## 3. RESULTS

METE’s *Ñ*_0_ was positively related to habitat cover, both immediately after habitat loss and after stabilization of community dynamics (Fig. 3, left panels). After stabilization, spatial clustering also affected *Ñ*_0_ to some extent. *Ñ*_0_ was highest under high habitat cover and low spatial clustering of habitat destruction.

**Figure 3.**
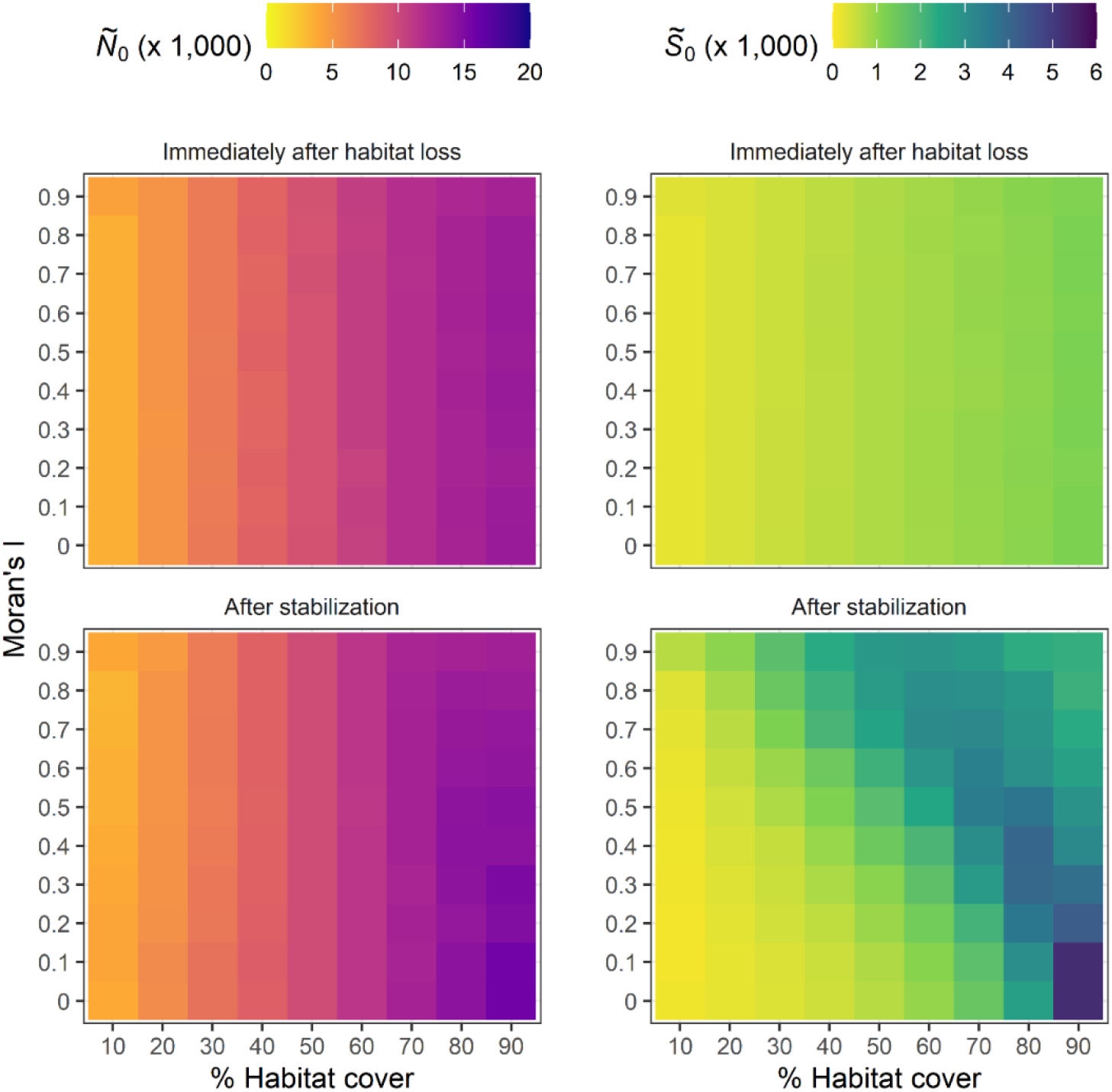
The Ñ_0_ and 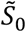 values that provided the best fitting METE approximation of the SAR observed in simulated data, per degree of spatial habitat aggregation (Moran’s I) and percentage habitat cover, immediately after habitat loss has occurred (top panels), and after subcommunity dynamics have stabilized (bottom panels).

Immediately after habitat loss, METE’s 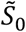 was positively related to habitat cover (Fig. 3, top right panel). Spatial clustering (Moran’s I) had no effect on 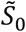. Yet, the long-term effects of habitat loss on METE’s 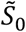 were less straightforward and depended on the interaction between habitat cover and spatial clustering (Fig. 3, bottom right panel). Highest levels of 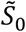 were attained at high habitat cover and high fragmentation levels, similar to Fahrig (2017). Here, small gaps in the continuous landscape caused a more diverse habitat, which provided niches for additional species. With decreasing habitat cover, the peak in the 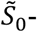-curve shifts towards high spatial clustering of habitat patches.

Empirically derived SARs are best represented by METE’s estimation of SARs immediately after habitat loss (Fig. 4). In all cases, species richness was much lower than what it should be if the areas had remained untouched by humans. Results from our model simulations show, however, that species richness may improve over time, especially in areas where forest cover has remained relatively high.

**Figure 4.**
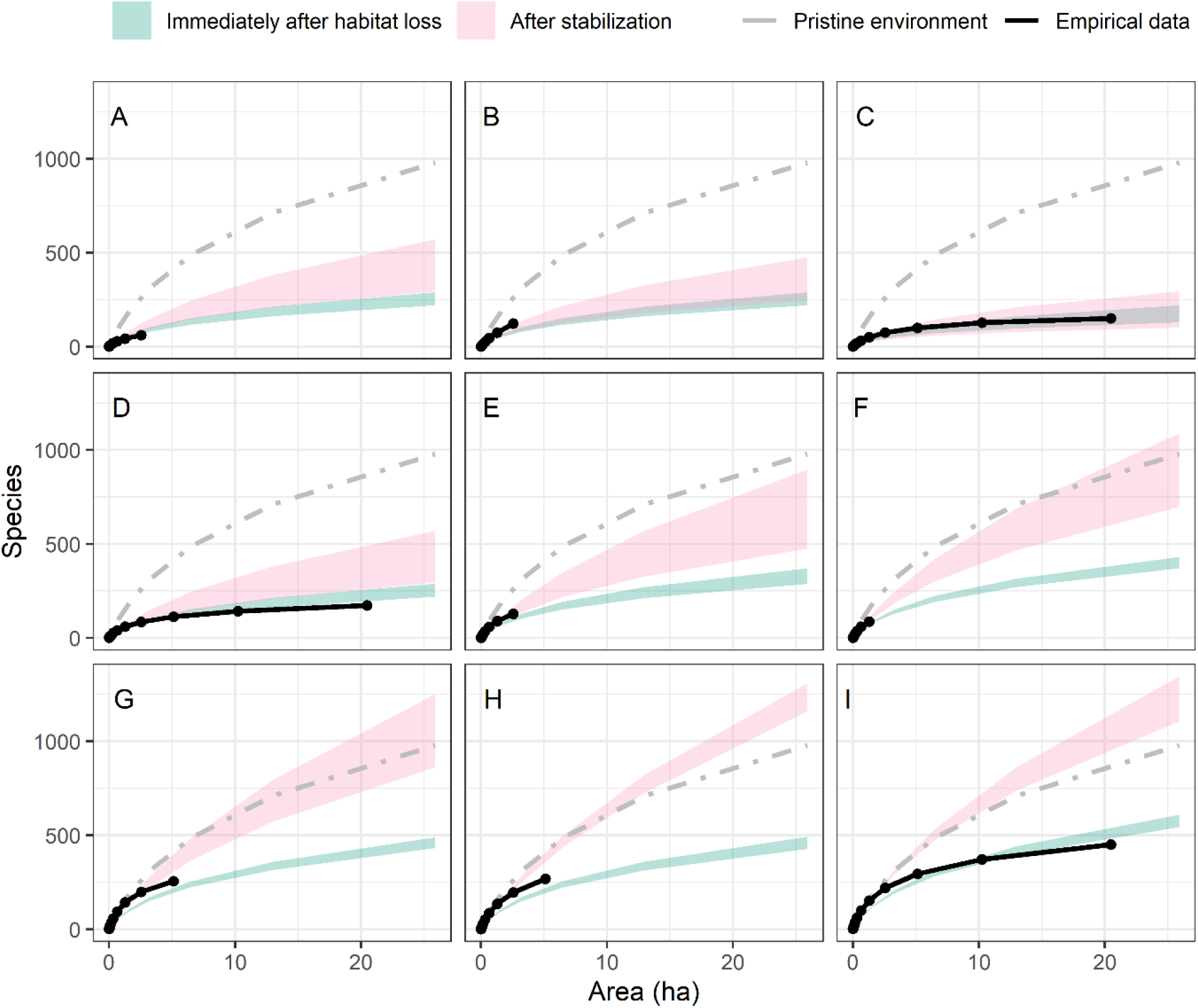
Species-Area Relationships per study area. Solid black points and lines indicate SARs generated using the empirical data. Dashed grey lines show the METE estimation of the SAR of the simulated pristine environment. Shaded regions represent the SAR-areas predicted with the adjusted METE, using the observed habitat cover and habitat fragmentation levels ± 5%, immediately after habitat loss (green) and after stabilization of community dynamics (pink), respectively. (**A, E, F**) Safar et al. (2022), (**B, D, G, I**) Pinho et al. (2021), (**C**) Santos et al. (2007), and (**H**) Magnago et al. (2015).

## 4. DISCUSSION

We demonstrate that species-area relationship (SAR) can be estimated in environments undergoing habitat loss and fragmentation, using METE predictions with its state variables S_0_ and N_0_ adjusted to the values corresponding to the environments’ degrees of habitat loss and fragmentation.

Our simulations produced some counterintuitive results. Under severe habitat loss and fragmentation, species richness continued to decline over time. However, when little habitat was lost, or when habitat loss was highly clustered and fragmentation was low, landscape-level species richness was higher after community dynamics had stabilized than immediately after habitat loss. This increase probably resulted from continued immigration from the metacommunity and the recolonization of areas from which species had been temporarily lost. Because species differed in their dispersal capacities, recolonization occurred at different rates among species and patches. Stochastic differences in colonization success may also have increased compositional variation among patches, thereby increasing landscape-level richness through greater beta diversity. Thus, although local richness may not have increased in every patch, the set of species present across the landscape became more diverse over time. These results differ from those of strictly neutral models, which assume equal dispersal capacities among species and therefore do not capture this source of variation in recolonization dynamics (Thompson et al. 2019). Especially in environments where forest cover levels remain high, many species continue to be sufficiently abundant, which reduces the risk of stochastic events leading to their local extinction. Meanwhile, the created gaps in habitat cause community dynamics to change. In the pristine, continuous landscape, long-distance dispersing species are at an advantage and are therefore overabundant. Yet, when gaps are introduced, locally dispersing species become more frequent, simply because they lose fewer seeds to the uninhabitable matrix. In environments where forest cover remains high but small gaps are present, a mixture of species with different dispersal capacities can exist. In addition, since species often show clustered distributions, the loss of habitat creates a mosaic of species diversity. If connectivity remains sufficient, this can maintain species diversity on a regional scale and can even increase species diversity locally. Furthermore, gaps generate edge effects, where species with niches other than those of the forest-center species can live and thrive. However, we did not account for differences in niches in our model.

While our model provides valuable insights into the impacts of habitat loss and fragmentation, its results should be interpreted with caution. A key assumption of our approach is that the metacommunity species pool remains intact in all scenarios we tested. This means our model applies specifically to areas where all species retain the ability to disperse into the focal area. If the focal area is located on the edge of the metacommunity or exists in isolation, the outcomes may differ significantly. In such cases, the dynamics of species loss and ecosystem responses could be fundamentally altered, and our model may not accurately reflect these conditions. Further research is needed to explore these edge cases and isolated systems, as they may reveal critical insights into the resilience of ecosystems under habitat loss.

In our model, we assume equal numbers of species with different dispersal capacities to be present in the metacommunity. As most Amazonian tree species are dispersed by means of zoochory, there are likely fewer locally dispersing species than we have assumed. If there indeed are fewer locally dispersing species within the metacommunity, our estimations of the Species-Area Relationships after stabilization of community dynamics are overshooting the actual SAR and should be adjusted to be closer to the SAR immediately after habitat loss.

The empirically derived SARs correspond most with the SARs estimated with METE immediately after habitat loss. This result indicates that community dynamics in these areas have not yet stabilized and are still ongoing. Immediately after habitat loss, fragmentation has no effect on species richness as the remaining species have not yet had the opportunity to respond to changes in their environment. Over time, however, fragmentation does play a significant role. In highly fragmented areas, subcommunities will diverge much more from one another in species composition than in contiguous areas. Many species will go extinct due to the stochastic processes occurring in such small subcommunities. In cases of high habitat cover and small gaps within the contiguous area (which is particularly prevalent in environments with low Moran’s I), both short-distance and long-distance dispersing species are able to coexist, resulting in an increased species richness. If our estimations are correct, species richness should increase in the two study areas with the highest remaining forest cover (Fig. 4H, I).

With this study, we are the first to provide objectively predicted species-area relationships in environments undergoing habitat loss and fragmentation, using an adjusted version of METE. Our empirical validation shows that we can successfully predict the loss in species richness following habitat loss. However, this should be considered a first attempt at adjusting METE. Future studies may focus on improving our method. For example, one could try to add the effects of spatial heterogeneity as a constraint on the Spatial Structure Function. Our current method, however, can already be used to predict changes in species richness due to habitat loss and fragmentation, thereby informing management practitioners of the tremendous impact of habitat destruction on biodiversity loss.

## AUTHOR CONTRIBUTIONS

**Monique de Jager:** Conceptualization, Analysis, Methodology, Visualization, Writing-Original draft, Writing-review & editing. **Finn J. van Wordragen:** Analysis, Investigation, Writing-review & editing. **Fleur Vrouwenvelder-Slegers:** Validation, Writing-Original draft, Writing-review & editing. **Edwin T. Pos:** Conceptualization, Methodology, Supervision, Writing-Original draft, Writing-review & editing.

## REFERENCES

Brummer, Alexander B., and Erica A. Newman. 2019. “Derivations of the Core Functions of the Maximum Entropy Theory of Ecology.” Entropy 21 (7): 712. 10.3390/e21070712.

Brush, Micah, Thomas J. Matthews, Paulo A. V. Borges, and John Harte. 2022. “Land Use Change through the Lens of Macroecology: Insights from Azorean Arthropods and the Maximum Entropy Theory of Ecology.” Ecography 2022 (8). 10.1111/ecog.06141.

Campos, Paulo R. A., Elias D. C. Neto, Viviane M. de Oliveira, and M. a. F. Gomes. 2012. “Neutral Communities in Fragmented Landscapes.” Oikos (Hoboken) 121 (11): 1737–48. 10.1111/j.1600-0706.2011.20336.x.

Campos, Paulo R. A., Alexandre Rosas, Viviane M. de Oliveira, and Marcelo A. F. Gomes. 2013. “Effect of Landscape Structure on Species Diversity.” PLOS ONE 8 (6): e66495. 10.1371/journal.pone.0066495.

Carey, Susan, John Harte, and Roger Del Moral. 2006. “Effect of Community Assembly and Primary Succession on the Species-Area Relationship in Disturbed Ecosystems.” Ecography 29 (6): 866–72. 10.1111/j.2006.0906-7590.04712.x.

Chisholm, Ryan A., Felix Lim, Yi Shuen Yeoh, Wei Wei Seah, Richard Condit, and James Rosindell. 2018. “Species-Area Relationships and Biodiversity Loss in Fragmented Landscapes.” Ecology Letters (Hoboken) 21 (6): 804–13. 10.1111/ele.12943.

Connor, Edward F., and Earl D. McCoy. 1979. “The Statistics and Biology of the Species-Area Relationship.” The American Naturalist 113 (6): 791–833. 10.1086/283438.

Diamond, Jm. 1989. “The Present, Past and Future of Human-Caused Extinctions.” Philosophical Transactions of the Royal Society of Londen Series B-Biological Sciences (London) 325 (1228): 469–77. 10.1098/rstb.1989.0100.

Diffenbaugh, Noah S., and Christopher B. Field. 2013. “Changes in Ecologically Critical Terrestrial Climate Conditions.” Science (Washington) 341 (6145): 486–92. 10.1126/science.1237123.

Fahrig, L. 2003. “Effects of Habitat Fragmentation on Biodiversity.” Annual Review of Ecology, Evolution, and Systematics (Palo Alto) 34: 487–515. 10.1146/annurev.ecolsys.34.011802.132419.

Fahrig, Lenore, Victor Arroyo-Rodriguez, Joseph R. Bennett, et al. 2019. “Is Habitat Fragmentation Bad for Biodiversity?” Biological Conservation (Oxford) 230 (February): 179–86. 10.1016/j.biocon.2018.12.026.

Franzman, Juliette, Micah Brush, Kaito Umemura, Courtenay A. Ray, Benjamin Blonder, and John Harte. 2021. “Shifting Macroecological Patterns and Static Theory Failure in a Stressed Alpine Plant Community.” Ecosphere 12 (6). 10.1002/ecs2.3548.

Fundacao, S. O. S. 2022. Atlas Dos Remanescentes Florestais Da Mata Atlântica, Relatório Técnico: Período 2020-2021. INPE.

Gonçalves-Souza, Thiago, Maurício Humberto Vancine, Nathan J. Sanders, et al. 2025. “LandFrag: A Dataset to Investigate the Effects of Forest Loss and Fragmentation on Biodiversity.” Global Ecology and Biogeography 34 (2). 10.1111/geb.70015.

Gorelick, Noel, M. Hancher, Mike Dixon, Simon Ilyushchenko, David Thau, and Rebecca Moore. 2017. “Google Earth Engine: Planetary-Scale Geospatial Analysis for Everyone.” Remote Sensing of Environment 202 (July): 18–27. 10.1016/j.rse.2017.06.031.

Haddad, Nick M., Lars A. Brudvig, Jean Clobert, et al. 2015. “Habitat Fragmentation and Its Lasting Impact on Earth’s Ecosystems.” Science Advances (Washington) 1 (2): e1500052. 10.1126/sciadv.1500052.

Hanski, Ilkka, Gustavo A. Zurita, M. Isabel Bellocq, and Joel Rybicki. 2013. “Species-Fragmented Area Relationship.” Proceedings of the National Academy of Sciences of the United States of America (Washington) 110 (31): 12715–20. 10.1073/pnas.1311491110.

Harte, J., T. Zillio, E. Conlisk, and A. B. Smith. 2008. “Maximum Entropy and the State-Variable Approach to Macroecology.” Ecology (Hoboken) 89 (10): 2700–2711. 10.1890/07-1369.1.

Harte, John. 2011. Maximum Entropy and Ecology: A Theory of Abundance, Distribution, and Energetics. Oxford University Press.

Harte, John, and Erica A. Newman. 2014. “Maximum Information Entropy: A Foundation for Ecological Theory.” Trends in Ecology & Evolution (London) 29 (7): 384–89. 10.1016/j.tree.2014.04.009.

Hubbell, Stephen P., Fangliang He, Richard Condit, Luís Borda-de-Água, James Kellner, and Hans ter Steege. 2008. “How Many Tree Species Are There in the Amazon and How Many of Them Will Go Extinct?” Proceedings of the National Academy of Sciences 105 (supplement_1): 11498–504. 10.1073/pnas.0801915105.

IPBES. 2019. Global Assessment Report on Biodiversity and Ecosystem Services of the Intergovernmental Science-Policy Platform on Biodiversity and Ecosystem Services. Zenodo. 10.5281/zenodo.6417333.

Jager, Monique de, and Edwin Pos. 2026. “Variation in Species’ Dispersal Capacities Amplifies Effects of Habitat Loss and Fragmentation on Biodiversity Loss.” Ecosphere 17 (1). 10.1002/ecs2.70516.

Keil, Petr, David Storch, and Walter Jetz. 2015. “On the Decline of Biodiversity Due to Area Loss.” Nature Communications (London) 6 (November): 8837. 10.1038/ncomms9837.

Kempton, R. A., and L. R. Taylor. 1974. “Log-Series and Log-Normal Parameters as Diversity Discriminants for the Lepidoptera.” Journal of Animal Ecology 43 (2): 381–99. 10.2307/3371.

Kuipers, Koen J. J., Roel May, and Francesca Verones. 2021. “Considering Habitat Conversion and Fragmentation in Characterisation Factors for Land-Use Impacts on Vertebrate Species Richness.” Science of The Total Environment 801 (December): 149737. 10.1016/j.scitotenv.2021.149737.

Larrey-Lassalle, Pyrène, Antoine Esnouf, Philippe Roux, Miguel Lopez-Ferber, Ralph K. Rosenbaum, and Eléonore Loiseau. 2018. “A Methodology to Assess Habitat Fragmentation Effects through Regional Indexes: Illustration with Forest Biodiversity Hotspots.” Ecological Indicators 89 (June): 543–51. 10.1016/j.ecolind.2018.01.068.

Larrey-Lassalle, Pyrène, Eléonore Loiseau, Philippe Roux, Miguel Lopez-Ferber, and Ralph K. Rosenbaum. 2018. “Developing Characterisation Factors for Land Fragmentation Impacts on Biodiversity in LCA: Key Learnings from a Sugarcane Case Study.” The International Journal of Life Cycle Assessment 23 (11): 2126–36. 10.1007/s11367-018-1449-5.

Lenore Fahrig. 2017. “Ecological Responses to Habitat Fragmentation Per Se.” Annual Review of Ecology, Evolution, and Systematics 48: 1–23. 10.1146/annurev-ecolsys-110316-022612.

Liang, Jingjing, Thomas W. Crowther, Nicolas Picard, et al. 2016. “Positive Biodiversity-Productivity Relationship Predominant in Global Forests.” Science (Washington) 354 (6309): aaf8957. 10.1126/science.aaf8957.

Loreau, M., S. Naeem, P. Inchausti, et al. 2001. “Ecology - Biodiversity and Ecosystem Functioning: Current Knowledge and Future Challenges.” Science (Washington) 294 (5543): 804–8. 10.1126/science.1064088.

Magnago, Luiz Fernando Silva, Ainhoa Magrach, William F. Laurance, et al. 2015. “Would Protecting Tropical Forest Fragments Provide Carbon and Biodiversity Cobenefits under REDD+?” Global Change Biology 21 (9): 3455–68. 10.1111/gcb.12937.

May, Felix, Benjamin Rosenbaum, Frank M. Schurr, and Jonathan M. Chase. 2019. “The Geometry of Habitat Fragmentation: Effects of Species Distribution Patterns on Extinction Risk Due to Habitat Conversion.” Ecology and Evolution (Hoboken) 9 (5): 2775–90. 10.1002/ece3.4951.

Nath, Debashis, Reshmita Nath, and Wen Chen. 2024. “Faster Dieback of Rainforests Altering Tropical Carbon Sinks under Climate Change.” Climate and Atmospheric Science 7 (1): 235. 10.1038/s41612-024-00793-0.

Newman, Erica A., Mark Q. Wilber, Karen E. Kopper, et al. 2020. “Disturbance Macroecology: A Comparative Study of Community Structure Metrics in a High-Severity Disturbance Regime.” Ecosphere (Hoboken) 11 (1). 10.1002/ecs2.3022.

Pereira, Henrique M., and Gretchen C. Daily. 2006. “Modeling Biodiversity Dynamics in Countryside Landscapes.” Ecology 87 (8): 1877–85. 10.1890/0012-9658(2006)87[1877:MBDICL]2.0.CO;2.

Pinho, Bruno X., Marcelo Tabarelli, Cajo J. F. ter Braak, et al. 2021. “Functional Biogeography of Neotropical Moist Forests: Trait–Climate Relationships and Assembly Patterns of Tree Communities.” Global Ecology and Biogeography 30 (7): 1430–46. 10.1111/geb.13309.

Quinn, Jf, and Sp Harrison. 1988. “Effects of Habitat Fragmentation and Isolation on Species Richness - Evidence from Biogeographic Patterns.” Oecologia (New York) 75 (1): 132–40. 10.1007/BF00378826.

Reich, Peter B., David Tilman, Forest Isbell, et al. 2012. “Impacts of Biodiversity Loss Escalate Through Time as Redundancy Fades.” Science (Washington) 336 (6081): 589–92. 10.1126/science.1217909.

Revenga, Carmen, Jake Brunner, Norbert Henninger, Ken Kassem, and Richard Payne. 2000. “Pilot Analysis of Global Ecosystems: Freshwater Systems.” World Resources Institute.

Ricketts, Taylor H. 1999. Terrestrial Ecoregions of North America: A Conservation Assessment. Island Press.

Rominger, A. J., K. R. Goodman, J. Y. Lim, et al. 2016. “Community Assembly on Isolated Islands: Macroecology Meets Evolution.” Global Ecology and Biogeography 25 (7): 769–80. 10.1111/geb.12341.

Rominger, Andrew J., and Cory Merow. 2017. “meteR: An r Package for Testing the Maximum Entropy Theory of Ecology.” Methods in Ecology and Evolution 8 (2): 241–47. 10.1111/2041-210X.12625.

Safar, Nathalia V. H., Masha van der Sande, Carlos Ernesto G. R. Schaefer, et al. 2022. “Landscape Openness Has Different Effects on the Structure, Diversity and Functional Composition of Brazilian Rainforests.” Forest Ecology and Management (Amsterdam) 520 (September): 120395. 10.1016/j.foreco.2022.120395.

Santos, Bráulio Almeida, Carlos A. Peres, Marcondes Albuquerque de Oliveira, Alexandre S. Grillo, Cecília P. Alves-Costa, and Marcelo Tabarelli. 2007. “Drastic Erosion in Functional Attributes of Tree Assemblages in Atlantic Forest Fragments of Northeastern Brazil.” Biological Conservation 141 (1): 249–60. 10.1016/j.biocon.2007.09.018.

Seibold, Sebastian, Claus Baessler, Roland Brandl, et al. 2017. “An Experimental Test of the Habitat-Amount Hypothesis for Saproxylic Beetles in a Forested Region.” Ecology (Hoboken) 98 (6): 1613–22. 10.1002/ecy.1819.

Steege, H. ter, N. Pitman, D. Sabatier, et al. 2003. “A Spatial Model of Tree α-Diversity and Tree Density for the Amazon.” Biodiversity and Conservation (Dordrecht) 12 (11): 2255–77. 10.1023/A:1024593414624.

Steege, Hans ter, Paulo I. Prado, Renato A. F. de Lima, et al. 2020. “Biased-Corrected Richness Estimates for the Amazonian Tree Flora.” Scientific Reports (Berlin) 10 (1): 10130. 10.1038/s41598-020-66686-3.

Steege, Hans ter, Daniel Sabatier, Sylvia Mota de Oliveira, et al. 2017. “Estimating Species Richness in Hyper-Diverse Large Tree Communities.” Ecology 98 (5): 1444–54. 10.1002/ecy.1813.

Supp, S. R., X. Xiao, S. K. M. Ernest, and E. P. White. 2012. “An Experimental Test of the Response of Macroecological Patterns to Altered Species Interactions.” Ecology 93 (12): 2505–11. 10.1890/12-0370.1.

Taubert, Franziska, Rico Fischer, Juergen Groeneveld, et al. 2018. “Global Patterns of Tropical Forest Fragmentation.” Nature (Berlin) 554 (7693): 519–+. 10.1038/nature25508.

Thompson, Samuel E. D., Ryan A. Chisholm, and James Rosindell. 2019. “Characterising Extinction Debt Following Habitat Fragmentation Using Neutral Theory.” Ecology Letters (Hoboken) 22 (12): 2087–96. 10.1111/ele.13398.

Tilman, D., C. L. Lehman, and K. T. Thomson. 1997. “Plant Diversity and Ecosystem Productivity: Theoretical Considerations.” Proceedings of the National Academy of Sciences of the United States of America (Washington) 94 (5): 1857–61. 10.1073/pnas.94.5.1857.

Tscharntke, T., I. Steffan-Dewenter, A. Kruess, and C. Thies. 2002. “Contribution of Small Habitat Fragments to Conservation of Insect Communities of Grassland-Cropland Landscapes.” Ecological Applications (Hoboken) 12 (2): 354–63. 10.1890/1051-0761(2002)012[0354:COSHFT]2.0.CO;2.

